# Graph Lens Lite: A browser-based tool for interactive visualization and exploration of biological networks

**DOI:** 10.64898/2026.02.25.708026

**Authors:** Matthias Ley, Kinga Kęska-Izworska, Lucas Fillinger, Samuel M Walter, Fabio Baumgärtel, Enrico Bono, Louiza Galou, Peter Andorfer, Pirmin Hauser, Johannes Leierer, Klaus Kratochwill, Paul Perco

## Abstract

Biological network visualization together with graph-based analyses are key techniques in systems biology and network medicine to detect patterns and generate hypotheses regarding disease pathobiology, drug target identification, biomarker prioritization, and digital drug discovery. Network representations provide an intuitive way to communicate and share research findings. We have developed Graph Lens Lite, a browser-based tool that combines rich visualization with a streamlined interface for exploring and sharing biological networks. It offers an expressive query language, topological network analysis, interactive filtering, visual grouping, customizable layouts, a data editor, fine-grained property-based styling, animated edge-flow visualization, community detection, and a context-aware locally powered AI assistant particularly suited for exploring molecular models of disease pathobiology or drug mechanism of action. We demonstrate its utility on a curated network model of autosomal dominant polycystic kidney disease. Graph Lens Lite is open source, with a live web version available at https://delta4ai.github.io/GraphLensLite/.

## 1 Introduction

Visualization of biological networks is a core technique in systems biology for representation and analysis of complex, often heterogeneous, data. Network visualization combined with quantitative graph analysis such as detection of network modules, hub or bottleneck nodes, or cross-talk between networks can lead to novel hypothesis generation or detection of relevant patterns in biomedical data (1). Biological networks are of particular relevance in digital drug discovery and drug target prioritization and are used in modeling disease pathobiology as well as drug mechanism of action on a molecular level. Biological networks also represent an intuitive way of communicating and sharing research findings with collaborators or disseminating them to the broader scientific community.

Cytoscape (2) and Gephi (3) are considered gold standards for interactive network visualization with an exhaustive plugin ecosystem and rich layout capabilities, but are desktop-focused and lack portability or modern web-app features. Other tools such as STRING (4) or OmicsNet 2.0 (5) prioritize data integration, enrichment analysis, and access to extensive biological databases over visualization and customization, integrating the network viewer as a helper and not as the focus of the application. Several web-based alternatives have emerged to enhance portability. Cytoscape Web (6) extends the Cytoscape ecosystem with collaborative workflows for network visualization, though focusing on core functionalities with basic filtering and styling. Gephi Lite (https://lite.gephi.org/) uses sigma.js (https://www.sigmajs.org/) for efficient WebGL-based rendering, demonstrating the viability of the engine for browser-based exploration, though it compromises customization, filtering, and advanced styling. Similarly, Graphia(7) provides both a desktop and web client with a powerful 2D/3D rendering engine, but focuses less on user-friendly styling, highlighting, grouping and filtering, prioritizing efficiency, enrichment, and layout.

We have developed Graph Lens Lite (GLL) to address the need for a browser-based tool that combines visualization power with an intuitive, streamlined interface, enabling researchers to rapidly explore and evaluate complex networks without requiring expertise in specialized software. GLL offers an expressive query language alongside graphical user interface (GUI)-based filtering, visual grouping, and finegrained styling controls. Users can compute network metrics, apply customizable layouts, and map data attributes to visual scales. GLL exports networks as portable JSON files that capture the full application state for seamless collaboration, or as data tables for downstream analysis.

## 2 Materials and methods

### 2.1 Implementation and architecture

We developed GLL using vanilla JavaScript with no front-end framework, bundled with esbuild (https://esbuild.github.io). For graph rendering and interaction, GLL uses sigma.js (https://www.sigmajs.org/), a WebGL-based rendering engine, with graphology (https://graphology.github.io/) as the underlying graph data model and for layout and metric computation. Custom WebGL shader modules extend the renderer with animated edge flows and a node-density heatmap layer. Spreadsheet import and export of node and edge tables use ExcelJS (https://github.com/exceljs/exceljs). The graph assistant renders Markdown responses with marked (https://marked.js.org/) and sanitizes the resulting HTML via whitelist approach using DOMPurify (https://github.com/cure53/DOMPurify). An in-memory cache store holds shared module states and the workspace data (coordinates, filters, styles, group memberships, active query and the syntax tree, and view states), which can be serialized to JSON for export, enabling session restoration. While the core application requires no internet access, two optional network paths exist: the STRING demo loader issues outbound requests to fetch example data, and the background service (see packaging and distribution) accepts inbound graph handoffs from external applications. No usage data leaves the client.

### 2.2 Query language

The query language is based on logical combinations of conditions on node and edge properties. Each clause pairs a property-path (“Section::Group::Property”) with a numeric range (“BETWEEN x AND y”), inverted range (“LOWER THAN x OR GREATER THAN y”), or categorical membership (“IN [..]”) operator. Clauses are combined with the connectors “AND”, “OR”, and “NOT”, support nested parentheses, and are evaluated left to right. The editor renders each token as spans with dedicated CSS classes for syntax highlighting, and validates on each change, flagging empty clauses, missing connectors, unmatched brackets, and invalid property paths. Valid queries are compiled into an abstract syntax tree that employs functions to evaluate each element to drive filtering and selection.

### 2.3 Graph assistant

The graph assistant’s system prompt encodes GLL’s main functionalities and instructs the model to reference UI controls using predefined glyphs that are rendered as clickable action buttons in its replies, and to request a query when the user asks to filter the graph. Query generation is split into two phases. In the first, on recognizing that a query is needed, the model appends a block (“«<QUERY_INTENT»>summary, scope«<END»>“) to its reply, which is stripped from the displayed response. In the second phase, a separate structured-output call (temperature 0) sends a JSON schema to the model using Ollama’s “format” parameter. The schema allows only the graph’s existing property paths as fields and only the query language’s operators. The model therefore cannot emit a non-existent property or an unsupported operator. The generated structured output is rendered to a query string that is then decoded and evaluated in the same manner as typed or GUI-built queries. If the structured output fails to render to a valid query (e.g. an empty membership list), a single repair pass retries, remapping any invalid property to its nearest valid path by Levenshtein distance.

### 2.4 Packaging and distribution

GLL is released under the MIT license. Desktop builds for Windows, macOS, and Linux are produced with electron-builder (https://www.electron.build/) on Electron 36 (https://www.electronjs.org/). A platform-independent single-file build is produced with inline-source (https://github.com/popeindustries/inline-source), which inlines the application bundle, stylesheets, and assets into one self-contained HTML file of about 2.6 MB that runs in any modern browser. Releases are versioned semantically and distributed through GitHub Releases and GitHub Pages, with a live demo at https://delta4ai.github.io/GraphLensLite/. GLL can also run as a background service built with Node.js (https://www.nodejs.org) that exposes a token-protected REST endpoint, accepting data in GLL’s JSON format and streaming it to connected browsers over Server-Sent Events, so external applications can hand off network data to GLL for interactive exploration, with concurrent users kept separate through named sessions, without writing intermediate files. The codebase is covered by over 1,300 automated tests across 66 files, written with Vitest (https://vitest.dev/).

## 3 Results

GLL presents network exploration through a single-page interface built around a central network view and a set of dedicated panels for data input, filtering, selection, styling, metrics, data and query editors, and an assistant (Figure 1).

**Figure 1:**
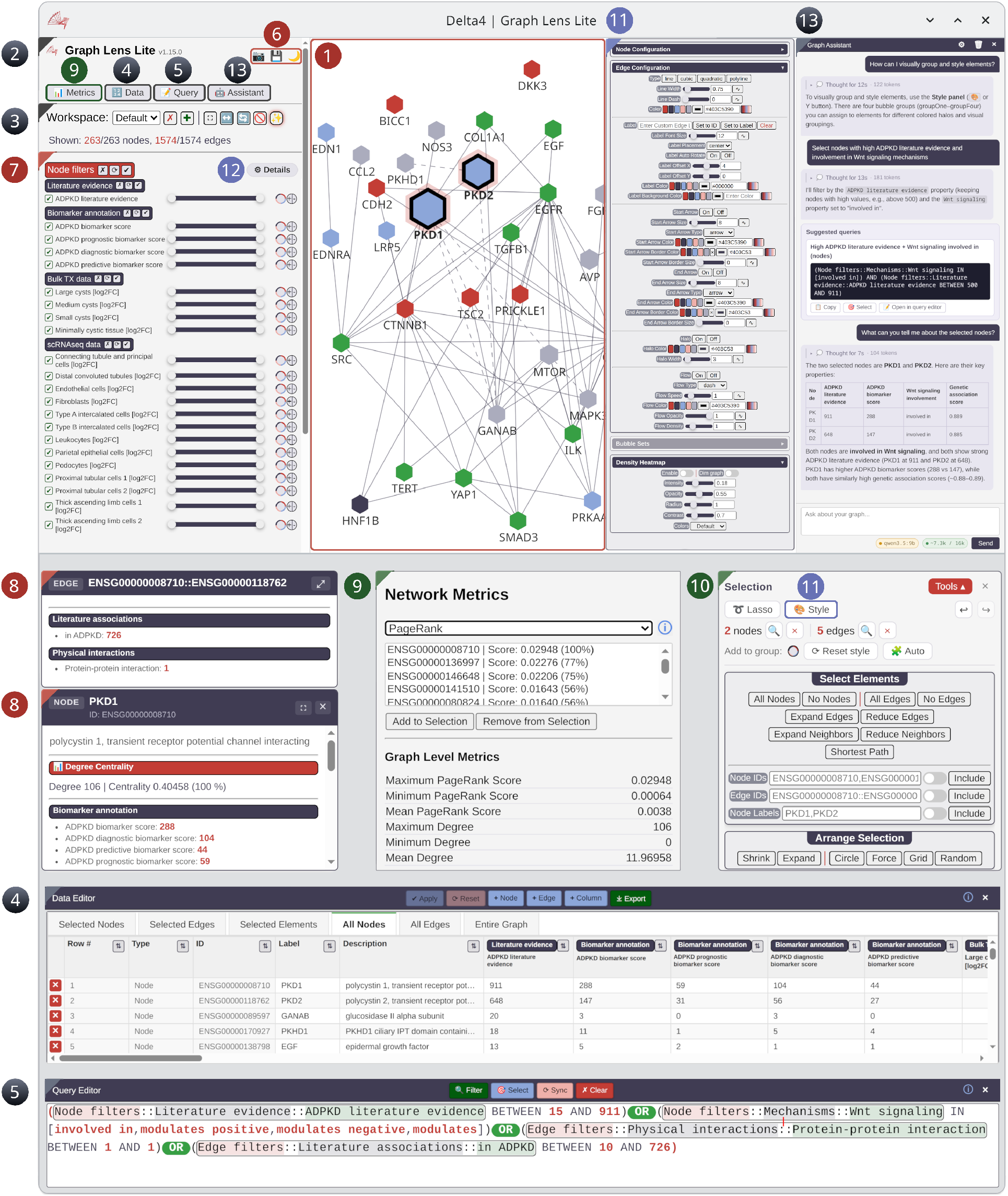
The graphical user interface of Graph Lens Lite. (1) Main network view, (2) File import, (3) Workspace management, (4) Data editor, (5) Query editor, (6) Network and image export, (7) Node and edge filtering, (8) Node and edge tooltips, (9) Network metrics, (10) Selection panel, (11) Style panel, (12) Edit mode, (13) Graph assistant.

### 3.1 Data input

GLL accepts spreadsheets containing node and edge tables, or a JSON format preserving the application state. Input data require an “ID” column for nodes and “Source ID/Target ID” columns for edges. Optional columns may specify visual attributes such as labels, shapes, sizes, colors, or coordinates. Custom properties containing user data can be added as new columns, with optional group labels in square brackets. A template with column specifications, example values, and supported data types is available on GitHub and can also be generated from within the application. Additionally, a demo loader enables direct fetching of protein-protein interaction networks from the STRING(4) database. Separately, a running GLL service instance can receive graphs programmatically from external applications over a token-protected endpoint (see packaging and distribution), without manual file handling.

### 3.2 Guided tour

On application launch, users can start an interactive guided tour that loads a small example protein network and walks the user step-by-step through the application’s core components. At each step the tour highlights relevant icons, opens panels, and explains the usage with live interface components, lowering the entry barrier for new users.

### 3.3 Graphical user interface

The numbered elements in Figure 1 correspond to the following components:

1. **Main network view:** The central window for displaying and interactively exploring the loaded network through panning, zooming, node dragging, hovering over elements to highlight adjacent elements, and an overview minimap.
2. **File import:** Open the launch screen to load network data from spreadsheets (node and edge tables) or portable GLL JSON files, or fetch protein-protein interaction data from the STRING database.
3. **Workspace management:** Create and switch between independent workspaces, each preserving its own layout, filters, queries, groupings, and styles. Reset the layout, fit the graph to view, hide disconnected nodes, or toggle highlight effects.
4. **Data editor:** Spreadsheet-style interactive table for adding, deleting, and editing nodes, edges, and their properties, defining new custom property columns, and export the entire network, nodes, edges, or filtered sub-networks.
5. **Query editor:** Editor for nested graph filtering and selection using a custom query language, supporting Boolean, comparison, and set-membership operators. Synchronizes with the GUI filtering panel.
6. **Network and image export:** Save the complete application state to a portable GLL JSON file for session restoration and sharing, or export the current view as high-resolution PNG.
7. **Node and edge filtering:** GUI-based selection and visual grouping, filtering via range sliders for numeric properties and category checklists for categorical values, synchronized with the query language.
8. **Node and edge tooltips:** Displays metadata for the selected node or edge element on click.
9. **Network metrics:** Compute topological centrality measures (degree, betweenness, closeness, eigenvector, and PageRank) to rank, filter, and select graph elements, with graph-level statistics and illustrative documentation for each method.
10. **Selection panel:** Shows current node and edge selection counts with lasso selection and undo/redo history. An expandable section provides additional focus, search, neighbor/edge expansion, shortest-path tracing between two selected nodes, and per-selection sub-layout controls.
11. **Style panel:** Edit node, edge, and bubble-group styles, and map data properties or computed network metrics onto visual scales (continuous and discrete color scales, numeric size scales).
12. **Edit mode:** Toggles fine-grained configuration for GUI-based filtering via precise numeric thresholds.
13. **Graph assistant:** Context-aware AI assistant that answers natural-language questions about the network, translates them into runnable queries, and summarizes the current selection.

### 3.4 Features and workflow

GLL starts by loading network data from spreadsheet files containing node and edge tables (Figure 1.2) or from JSON files. User-supplied properties (coordinates, styling options) are applied where specified. Alternatively, users can fetch demo data from the STRING database. The workspace management panel (Figure 1.3) allows switching between independent environments, each preserving its own aesthetics, layouts, filters, and queries.

**Figure 2:**
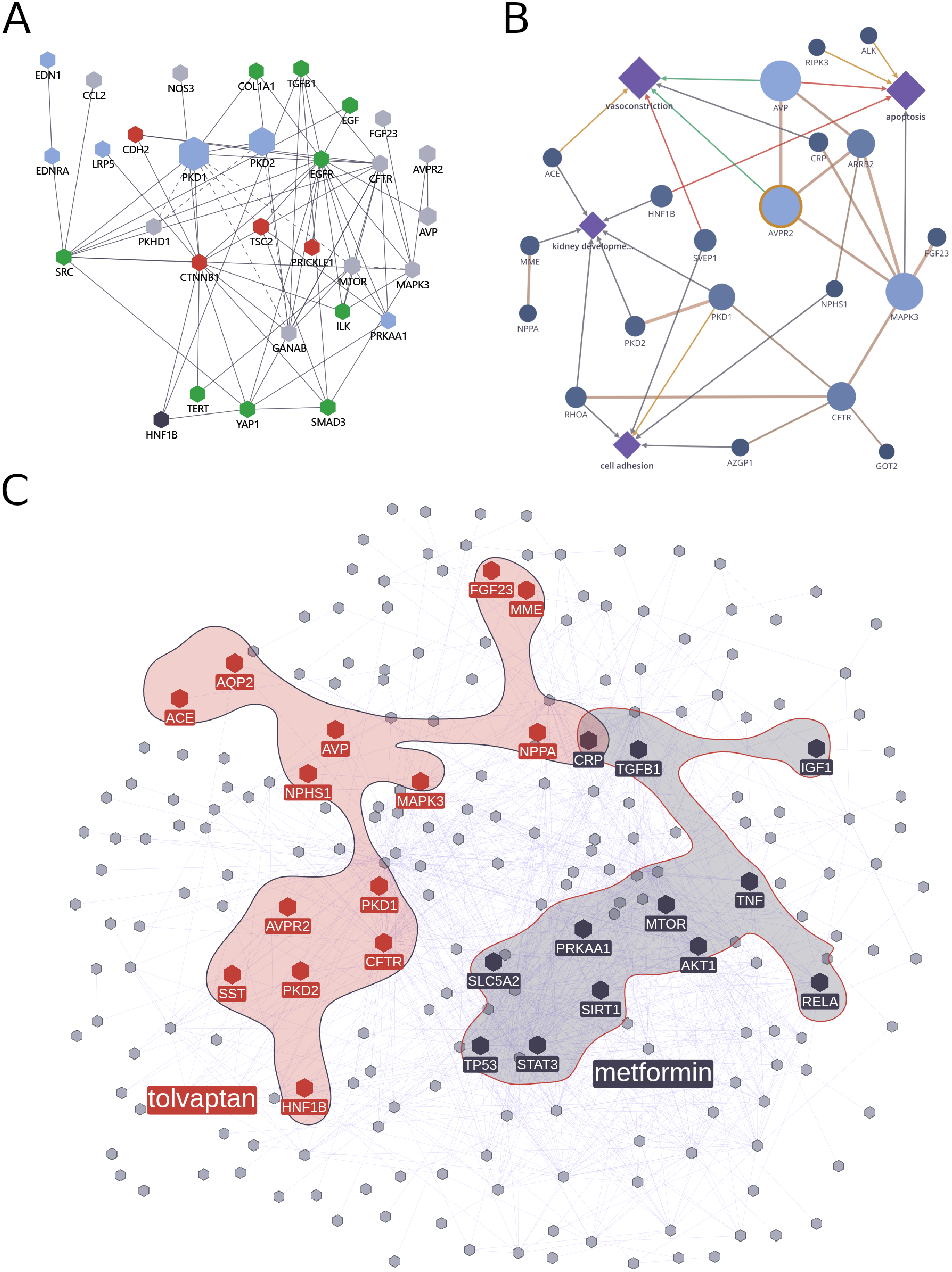
Visualizing disease and drug models. Panel A: Proteins of the ADPKD molecular model being associated to the Wnt signaling pathway (in green for positive modulation of the pathway, red for negative modulation of the pathway, dark blue for modulation of the pathway, and light blue for genes being pathway members without any further information on regulation) as well as additional molecules being associated with ADPKD that are linked to members of the Wnt signaling pathway (in gray) via either protein-protein interactions or co-annotations in the context of ADPKD. Panel B: The drug MoA molecular model for tolvaptan showing key associated proteins as well as associated molecular mechanisms. Panel C: The largest connected component of the ADPKD molecular model with interference signatures for tolvaptan and metformin given, indicating their complementary impact on ADPKD disease pathobiology.

Once loaded, users can explore the network structure through multiple approaches. The selection panel (Figure 1.10) enables element search, selection, expansion to connected neighbors, and tracing the shortest path between two selected nodes on the currently visible network. Metadata for selected node and edge elements appear in specific tooltips (Figure 1.8). For deeper analysis, GLL computes five node-centrality measures (degree (8), betweenness (8), closeness (9), eigenvector (10) and PageRank (11)) together with graph-level statistics including network density and centralization, to identify key nodes (Figure 1.9). Users can select a metric and examine one or more highly ranked nodes, with graph-level statistics displayed in the metric panel. In-app documentation describes each metric’s methodology and provides references to relevant literature. The network can be arranged with a range of layout algorithms: live-animating force-directed, hierarchical, circular, radial, concentric, and circle-packing, applied either to a selection or across the entire workspace, with animated transitions between layouts and an optional overlap-removal pass.

GLL supports network filtering through graphical controls (Figure 1.7), an edit mode for precise numeric input (Figure 1.12), and a query editor for complex, nested operations (Figure 1.5). All interfaces are synchronized, acting on the same underlying node and edge properties. The query language is a small, purpose-built domain-specific language (DSL) rather than a free-text search. The editor validates as the user types, highlighting empty clauses, missing connectors, and unmatched brackets. Validation is grounded in the property hierarchy. Valid queries compile to an expression tree evaluated per element, so that single statements drive both filtering and selection while staying consistent with the GUI controls. Contextual help is accessible via an adjacent icon.

GLL integrates an optional, locally hosted, context-aware AI assistant (Figure 1.13), lowering the barrier to exploration by translating questions into instructions, queries and summaries. Each request embeds a snapshot of the current workspace, including node and edge counts, the property hierarchy with data types, active filters, the current query, and the currently selected nodes and edges including their property values and computed network metrics. Responses are grounded in the live graph state. When asked for a query, the assistant constructs a schema-constrained structured query representation based on the loaded graph’s properties. The query is constrained to the data, referencing only real properties and supported operators. Results can be applied to select elements via rendered action buttons, or loaded into the query editor for further refinements.

The assistant communicates with an Ollama server and is local by default, with configuration handled in the panel’s settings. Enabling it requires a running Ollama installation and a downloaded instruction-following model capable of reliably creating structured JSON output. Response quality depends on the chosen model. Qwen3.5:9b produced strong results while running efficiently on consumer-grade graphics processing units (GPUs)(12). Running a local endpoint prevents transmission of potentially sensitive network data to third-party services. A context-budget indicator reports how much of the selected model’s context window the next request will consume and warns users when a request would exceed the threshold and be truncated.

The styling panel (Figure 1.11) offers extensive visual customization options for nodes, edges, and groups. Properties such as geometry (shape, size), color (fill, border, line), labels (text, placement, font), and annotations (badges, halos, arrows) can be configured independently for each element type. Edges can additionally render animated directional flow, with selectable flow styles (e.g. comet or chevron), adjustable opacity and density, to express directionality or magnitude. A node-density heatmap can be drawn to highlight dense regions of the layout, with controls for intensity, opacity, radius, contrast, thresholds, and predefined color maps. Up to four bubble sets allow visual grouping of nodes with adjustable appearance, populated manually or automatically through Louvain community detection. Numeric and categorical attributes, whether user-supplied or computed from network metrics, can be dynamically mapped to visual features like color and size to highlight importance. Nodes can be rendered as pie charts that encode several numeric or categorical properties as wedges, adding a data dimension beyond color and size.

GLL includes an integrated data editor (Figure 1.4) for modifying network data in tabular format. Supported operations include column sorting, cell editing, node and edge addition or deletion, table reset, and export to spreadsheet format. Users can also add new columns to define custom properties for nodes or edges, which become immediately available in the filtering and styling interfaces, enabling graph creation and manipulation entirely within the application.

A graph can be exported in GLL’s native JSON format or as a high-resolution image (Figure 1.6). JSON exports preserve the complete application state, including views, filters, coordinates, and styling, enabling session restoration and collaborative sharing.

### 3.5 Modeling autosomal dominant polycystic kidney disease

In this case study, we use GLL to visualize the molecular pathobiology of autosomal dominant polycystic kidney disease (ADPKD), two drugs, and the interference signatures that link each drug to the disease network. The corresponding datasets are provided as Excel files (https://github.com/Delta4AI/GraphLensLite/tree/main/manuscript) so readers can reproduce these views and explore the tool’s features interactively.

ADPKD is the most common inherited kidney disorder, characterized by the gradual development and enlargement of multiple cysts in both kidneys, which ultimately leads to a progressive deterioration in kidney function (13). ADPKD is a multi-factorial disease involving different disease mechanisms and molecular pathways. The exact mechanisms of ADPKD development and progression are still not fully understood and a better understanding of disease pathobiology is the basis for the discovery of novel drug targets and the development of new treatment options. We consolidated a set of biomedical data on ADPKD to model disease pathobiology and visualize molecular interactions using GLL. Literature associated ADPKD molecular features (genes and proteins) were extracted from Delta4’s Hyper-C software platform that consists of sentence-level literature co-annotations between individual molecular features and ADPKD. This set of literature-associated ADPKD molecular features was complemented by genes showing differential expression in ADPKD as compared to healthy tissue based on Omics data. We downloaded the ADPKD bulk RNA-Seq dataset with the accession number GSE7869 from the Gene Expression Omnibus and re-analyzed the dataset to generate lists of differentially expressed genes (14). Global gene expression profiling was conducted on renal cysts of different volumes, and non-cancerous renal cortical tissue from nephrectomized kidneys was used as controls. Differential gene expression analysis involved data normalization, statistical modeling using linear models, identification of regulated genes based on log_2_ fold-change (log_2_ FC) and adjusted p-values. Ultimately, the ADPKD phenotype model was enriched with 20 genes which showed absolute log_2_ FC > 2 and adjusted p-values < 0.05 across all four comparisons (large, medium, and small cysts, as well as minimally cystic tissue as compared to healthy tissue from nephrectomized kidneys respectively). In addition, we re-analyzed a single-nucleus RNA sequencing (sn-RNA-Seq) dataset with the GEO accession number GSE185948, offering insights into the cellular heterogeneity of ADPKD (15). Genes with a p-value < 0.05 were included as candidates in the analysis. Ultimately, the ADPKD phenotype molecular model was enriched with genes that were expressed in at least one cell type with absolute log_2_ FC values > 2.5. In addition we retrieved information on genetic associations with ADPKD from the Open-Targets platform (16). Expression in healthy tissue and relevant renal cell types of molecular features of the ADPKD molecular model, quantified as normalized transcripts per million (nTPMs), were extracted from the Human Protein Atlas consensus expression profile dataset(17). Information on subcellular location for features of the ADPKD molecular model was retrieved from UniProt (18). Finally, we assigned molecular features of the ADPKD molecular model to mechanisms being discussed in recent reviews to be associated with ADPKD disease development and progression. Assignment of molecular features to mechanisms was done using the Gene Ontology (GO) annotation.

The final constructed ADPKD molecular model consisted of 263 molecular features that were connected by 1,574 edges (**Supplementary datafile S1**). Edges in the constructed network consisted of direct protein-protein interactions consolidated from IntAct (19), BioGRID (20), and Reactome (21) as well as of literature sentence-level co-annotations of publications focusing on ADPKD using the catalogs of molecular features and phenotypes as given in Delta4’s Hyper-C software platform.

The constructed molecular model can now be used to explore the pathobiology of ADPKD by either filtering for (i) certain molecular mechanisms, (ii) individual cell types, (iii) the most up- or down-regulated genes in renal tissue or specific cell types, (iv) the most central genes based on graph properties like node degree or betweenness among others. Figure 2A for example shows genes of the Wnt signaling pathway (in green for positive modulation of the pathway, red for negative modulation of the pathway, dark blue for modulation of the pathway, and light blue for genes being pathway members without any further information on regulation) as well as additional molecules being associated with ADPKD that are linked to members of the Wnt signaling pathway (in gray) via either protein-protein interactions or co-annotations in the context of ADPKD. This network view allows investigating novel connections of Wnt signaling pathway members with ADPKD-associated proteins.

The molecular model can subsequently be used to (i) screen for compounds showing beneficial impact on dysregulated disease mechanisms as shown previously in other renal diseases (22), (ii) identify drug targets of interest, investigate cell-cell interaction analysis to identify relevant receptor-ligand pairs following previously reported analyses (23), or (iii) prioritize drug targets for the development of new chemical entities.

The only FDA-approved treatment for ADPKD is the selective vasopressin V2-receptor (AVPR2) antagonist tolvaptan. We generated a molecular mechanism of action model for tolvaptan based on literature associated tolvaptan molecular features (genes and proteins) (Figure 2B). Molecular features were extracted from Delta4’s Hyper-C software platform that consists of sentence-level literature co-annotations between individual molecular features and tolvaptan. Mechanisms were assigned to molecular features via GO annotation. The constructed MoA molecular model can subsequently be used to assess drug mechanism of action on disease pathobiology via network interference analysis with interfering nodes between the ADPKD molecular model and the tolvaptan drug MoA molecular model making up the interference signature. We can subsequently also evaluate and graphically display how two different drugs interfere with disease pathobiology on the molecular level. Figure 2C displays the interference signature of tolvaptan (in red) and metformin (in blue), a compound currently being tested in an ongoing ADPKD clinical trial (NCT04939935) and being discussed as a compound to be combined with tolvaptan to treat patients with ADPKD (24). Network visualization suggests a complementary impact of the two drugs on ADPKD disease pathobiology due to the non-overlapping interfering signatures between ADPKD and the two drug models.

## 4 Discussion

We have developed GLL, an interactive, browser-based network viewer for exploring and sharing biological networks through a streamlined interface. It combines an expressive query language with topological network analysis, interactive filtering, visual grouping, customizable layouts, a data editor, finegrained property-based styling, animated edge-flow visualization, community detection, and a context-aware, locally hosted AI assistant, supporting the visual exploration of molecular models of disease pathobiology or drug mechanism of action. GLL requires only a web browser for core functionality: it is distributed as a single self-contained HTML file, which can be combined with a network model into a compact archive of a few megabytes for sharing, while platform-specific Electron bundles provide native desktop applications across Windows, macOS, and Linux. This minimal footprint enables collaboration, long-term stability, and offline operation.

Table 1 compares GLL against biomedical network exploration tools across their deployment surface (web, desktop, or single-file HTML), offline usability versus server dependency, portability of the exploration state, editability of the underlying data, the presence of a query language for complex filtering, and the availability of an AI assistant.

**Table 1:**
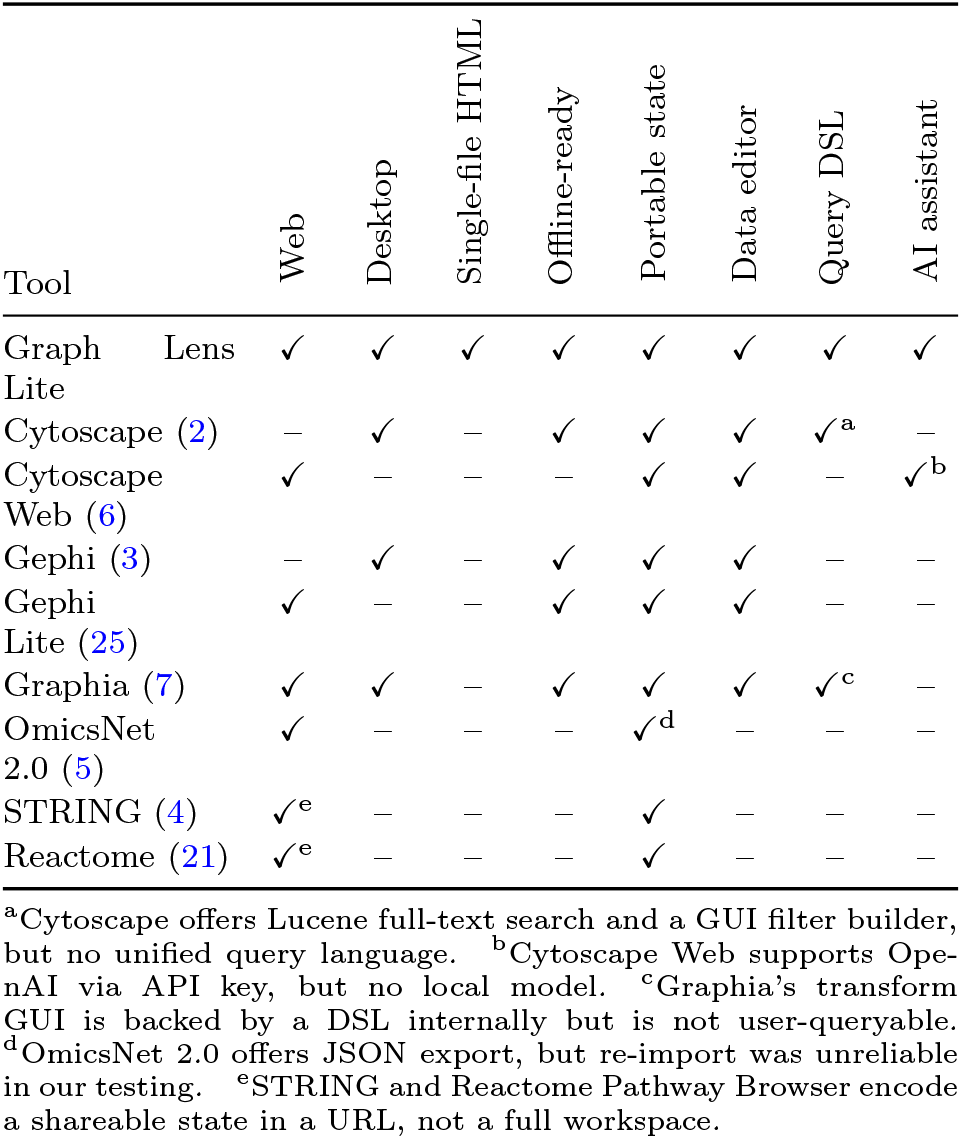
Comparison of selected network analysis and visualization tools for biomedical research. *Singlefile HTML*: the entire application is one HTML file that runs locally with no install procedure or server. *Offline-ready* : core functionality works without an external server or third-party API (optional demo data loaders may still require network access). *Portable state*: the complete application state (data, view, filters, layout, and styling) exports and reimports as a single file. *Data editor* : nodes, edges, and custom properties can be created, edited, and deleted from the GUI. *Query DSL*: a domain-specific query language for complex graph filtering is available. *AI assistant* : built-in assistant powered by a locally hosted open-source model for natural-language interaction. Cells: ✓ present, – absent.

While Table 1 provides a direct comparison across tools, some entries require further explanation to highlight GLL’s distinct capabilities. STRING Search(4) (https://string-db.org/cgi/input) and the Reactome Pathway Browser(21) (https://curator.reactome.org/PathwayBrowser) provide shareable states by encoding the current selection and analysis parameters into a URL, rather than serializing the full workspace. OmicsNet 2.0(5) offers JSON export whose re-import proved unreliable in our testing, reloading as a differently styled network including more than the exported set of nodes. To elaborate on domain specific query languages integrated into the application, Graphia’s(7) transform GUI is backed by a DSL internally but does not expose it for users to write or share, while Cytoscape(2) combines Lucene-style full-text search with a visual GUI-based filter engine, rather than a single, shareable query surface. GLL couples one validated query language with its GUI controls, so the same expression can be written by hand, constructed through the interface, or generated by the assistant. Of the web-based tools compared, only Cytoscape Web(6) provides a comparable assistant, but it routes requests to OpenAI (https://openai.com/) through a user-supplied API key with no local-model path, whereas GLL’s assistant runs against a locally hosted model so that graph data never leaves the machine. GLL therefore positions itself in a niche rather than replacing established platforms: it does not match the plugin ecosystems of Cytoscape and Gephi(3) or the database breadth of STRING, OmicsNet, and Reactome, but complements them with portable, offline-capable, and highly customizable exploration of focused, curated networks. Unlike canvas-based web renderers, GLL uses a GPU-accelerated WebGL engine, sustaining interactive exploration of large networks while retaining full styling, filtering, and grouping capabilities.

### 4.1 Limitations

GLL targets curated, interpretable networks rather than genome-scale topologies. WebGL rendering keeps panning, zooming, and selection interactive into the tens of thousands of elements, though larger networks combined with heavy per-element styling may still reduce responsiveness.

The network metric analysis covers five centrality measures computed on undirected graph data, and community structure can be revealed through Louvain community detection mapped onto visual groups. Directed-graph metrics are not yet supported. Path tracing between selected features is supported as an unweighted shortest path computed on the visible (filtered) subgraph. Weighted path costs are not considered, and only a single shortest path is returned when several of equal length exist.

While the current export/import flow reliably handles GLL’s JSON format, support for standardized network formats such as GraphML or CX2, and interoperability with established tools such as Cytoscape or Gephi could benefit applied biomedical researchers.

Keeping GLL’s query language and GUI synchronized proved difficult, as nested statements and negations do not map cleanly to graphical controls given the current design. The filtering panel therefore offers a single join mode applied across all clauses — OR by default or AND, the latter with an optional complete-cases mode — but cannot express the arbitrary nesting or negation supported by the raw query language. When a raw query is modified and applied to filter the graph, the GUI filtering panel is locked and flagged with a note that the query takes precedence, with the option to unlock it by resetting the query to match the GUI’s state. Value changes, property selections, and the join mode remain synchronized as the query is edited, whereas nesting and negation do not.

## Supporting information

Supplementary_Data_S1-ADPKD_model

## 5 Acknowledgements

The authors want to thank all working group members of the EU COST Action Program PerMediK (Personalized Medicine in Chronic Kidney Disease: Improved outcome based on Big Data) for fruitful discussions during the regular COST Action project meetings. Paul Perco and Matthias Ley are members of the EU COST Action Program PerMediK, CA21165.

## Author contributions

M.L.: conceptualization; resources; software; visualization; writing - original draft, review & editing. K.K.-I.: data curation; validation; writing - review & editing. L.F.: validation; writing - review & editing. S.W.: validation; writing - review & editing. F.B.: validation; writing - review & editing. E.B.: validation; writing - review & editing. L.G.: validation; writing - review & editing. P.A.: validation; writing - review & editing. P.H.: data curation; validation; writing - review & editing. J.L.: data curation; funding acquisition; validation; writing - review & editing. P.P.: conceptualization; data curation; funding acquisition; supervision; validation; writing - original draft, review & editing. K.K.: funding acquisition; supervision; validation; writing - review & editing.

## 8 Data availability

The application source code, distribution bundles, templates, the curated ADPKD molecular model, the drug mechanism of action model for tolvaptan, and the interference model are available at GitHub (https://github.com/Delta4AI/GraphLensLite) and Zenodo (https://doi.org/10.5281/zenodo.21035263). This website is free and open to all users and there is no login requirement.

